# LTBP1 promotes fibrillin incorporation into the extracellular matrix

**DOI:** 10.1101/2021.12.16.473056

**Authors:** Matthias Przyklenk, Veronika S. Georgieva, Fabian Metzen, Sebastian Mostert, Birgit Kobbe, Gerhard Sengle, Bent Brachvogel, Robert P. Mecham, Mats Paulsson, Raimund Wagener, Manuel Koch, Alvise Schiavinato

## Abstract

LTBP1 is a large extracellular matrix protein and an associated ligand of fibrillin-microfibrils. Knowledge of LTBP1 functions is largely limited to its role in targeting and sequestering TGFβ growth factors within the extracellular matrix, thereby regulating their bioavailability. However, the recent description of a wide spectrum of phenotypes in multiple tissues in patients harboring LTBP1 pathogenic variants suggests a multifaceted role of the protein in the homeostasis of connective tissues. To better understand the human pathology caused by LTBP1 deficiency it is important to investigate its functional role in extracellular matrix formation. In this study, we show that LTBP1 coordinates the incorporation of fibrillin-1 and −2 into the extracellular matrix *in vitro*. We also demonstrate that this function is differentially exerted by the two isoforms, the short and long forms of LTBP1. Thereby our findings uncover a novel TGFβ-independent LTBP1 function potentially contributing to the development of connective tissue disorders.

## Introduction

Fibrillin microfibrils are ubiquitous extracellular matrix (ECM) structures with essential functions during development and tissue homeostasis. Fibrillin-1 and fibrillin-2 form the core of “beads-on-a-string” microfibrils, to which an array of other proteins associate in a highly coordinated process. The resulting supramolecular networks represent multi-protein, multifunctional assemblies that show tissue-specific composition and function. Together with associated proteins, fibrillin microfibrils function as scaffolds for the correct deposition of elastin, and hence as necessary platforms for the formation of elastic fibers (1,2). Thus, fibrillins have a crucial role for conferring specific mechanical and structural properties to virtually all tissues in the organism (3). Fibrillin microfibrils are also involved in signaling by binding to several cellular receptors and by controlling the bioavailability of growth factors of the TGFβ family (4,5). The crucial role of these supramolecular structures for tissue homeostasis is highlighted by the vast and heterogeneous group of human genetic disorders that are commonly referred to as “micro/fibrillinopathies” (6,7). Among the large number of microfibril-associated proteins, the latent transforming growth factor beta binding proteins (LTBPs) play a particularly important role as both structural components and signaling hubs. They have a domain structure that is similar to that of fibrillins including tandem arrays of EGF like domains and interspersed unique “eight cysteine” or TB (TGFβ-binding proteinlike) domains (8). LTBPs are covalently linked to the propeptide of TGFβ and keep the cytokine in an inactive, latent conformation. LTBPs are encoded in mammals by four different genes (1–4). The third TB domains of LTBP1, −3 and −4 form two disulfide bonds with a pair of cysteines present in the latent associated peptide (LAP) of TGF□1, −2 and −3 to incorporate, store and eventually activate the latent cytokines within the ECM (9). LTBP1 is synthesized in two forms, a long (LTBP1L) and a short (LTBP1S) isoform, which are transcribed by alternative splicing from two distinct promoters (10,11). LTBP1L contains a stretch of 346 amino acid residues at its N-terminal region that is not present in the short form. Earlier studies have suggested that the long form is expressed during development and associates more efficiently with the ECM, while the short form is predominantly found in the adult (11,12).

Besides their function in regulating TGF□ signaling, experimental and clinical studies have revealed further roles of LTBPs as structural components of fibrillin microfibrils and elastic fibers. Studies in mice have shown that LTBP2 is necessary for the stability of microfibrils in the ciliary zonule (13,14). Nevertheless, it is not known how LTBP2 modulates the mechanical properties of fibrillin microfibrils. However, ectopic expression of LTBP4S in LTBP2-deficient mice could rescue the defects of the ciliary zonule. Furthermore *Ltbp2/4S* double knockout mice show more severe elastic fiber fragmentation together with more severe emphysematous changes in the lung and increased lethality when compared to *Ltbp4S* single knockout mice. Therefore, LTBP2 and LTBP4S have overlapping functions and may share similar mechanisms in the context of microfibril organization (15). LTBP4 is also essential for the linear deposition of fibulin-5 onto the elastic fibers in a process that is independent from its TGF□ binding activity (16).

Human mutations have been reported for all LTBPs, resulting in a spectrum of autosomal recessive Mendelian disorders unmistakably reflecting the diverse and partially overlapping functions of these proteins in virtually every tissue. Mutations in LTBP2, the only LTBP that does not bind to TGF□, have been linked to primary congenital glaucoma, ectopia lentis, megalocornea and Weill-Marchesani syndrome (17–19). LTBP3 pathogenic variants cause geleophysic dysplasia type 3 and platyspondyly with amelogenesis imperfecta (20,21). Mutations in LTBP4 cause cutis laxa type 1C (22). More recently pathogenic variants in *LTBP1* have been reported (23). Patients carrying LTBP1 truncating mutations show a complex spectrum of phenotypes including cutis laxa, craniofacial dysmorphism, mild cardiovascular defects and altered skeletal development and the term *LTBP1*-related cutis laxa syndrome has been proposed for these patients (23). Studies on patient-derived fibroblasts have suggested a complex pathomechanism involving both dysregulation of canonical TGF□ signaling and altered deposition of ECM components. However, even if specific roles in ECM assembly have been already described for LTBP2 (13-15,24) and LTBP4 (15,25,26), similar functions for LTBP1 have not yet been reported. In this study we show for the first time that LTBP1 has structural functions in the ECM, as it promotes the formation of fibrillin-1 and −2 microfibrils in cell cultures through a mechanism that appears to be independent from its function in regulating TGF□ signals. We also show that these functions are differentially fulfilled by LTBP1S and LTBP1L isoforms. This finding provides new information with clinical relevance, since it points to a novel pathomechanism for the newly identified *LTBP1*-related human disorders.

## Results

### LTBP1 promotes fibrillin-1 and −2 assembly downstream of fibronectin in mouse embryonic fibroblasts

Fibrillin-1 and −2 require fibronectin to be incorporated into the ECM (27,28). LTBP1 assembly also depends on fibronectin, but not on fibrillins (29,30). Therefore, assembly of fibrillins and LTBP1 can either occur independently, or LTBP1 may contribute to fibrillin incorporation downstream of fibronectin. To differentiate between these two alternatives, we transfected wild type (*Fn-WT*) mouse embryonic fibroblasts (MEFs) with a siRNA specific for *Ltbp1* and analyzed fibrillin-1 and −2 assembly by immunofluorescence. Strikingly, knockdown of *Ltbp1* resulted in a strong inhibition on the incorporation of both fibrillins, while having no effect on fibronectin (Figure 1A). qPCR analysis confirmed a 90% reduction of the *Ltbp1* mRNA level 5 days after transfection when compared to cells transfected with a control siRNA. Moreover, silencing of *Ltbp1* did not significantly affect the mRNA expression levels of *Fbn1* (fibrillin-1), *Fbn2* (fibrillin-2) or *Fn* (fibronectin) (Figure 1B). On the other hand, siRNA mediated double knockdown of both fibrillin-1 and −2 did not prevent LTBP1 or fibronectin incorporation into the ECM. As expected, fibronectin knockout MEFs (*Fn-KO*) failed to incorporate LTBP1, fibrillin-1 and fibrillin-2 into the ECM (Figure 1A). By performing qPCR analysis, we observed that fibronectin-deficient fibroblasts expressed *Ltbp1* at a significantly lower level than wild-type cells (Figure 1B). To verify that fibronectin is required for LTBP1 incorporation, we administered recombinant LTBP1S to fibronectin knockout and wild-type MEFs. While wild-type cells efficiently incorporated the exogenous protein into the ECM, fibronectin-deficient cells did not (Supplementary Figure 1). Therefore, fibronectin is required for the assembly of fibrillin microfibrils by supporting the incorporation of LTBP1 into the ECM.

**Figure 1.**
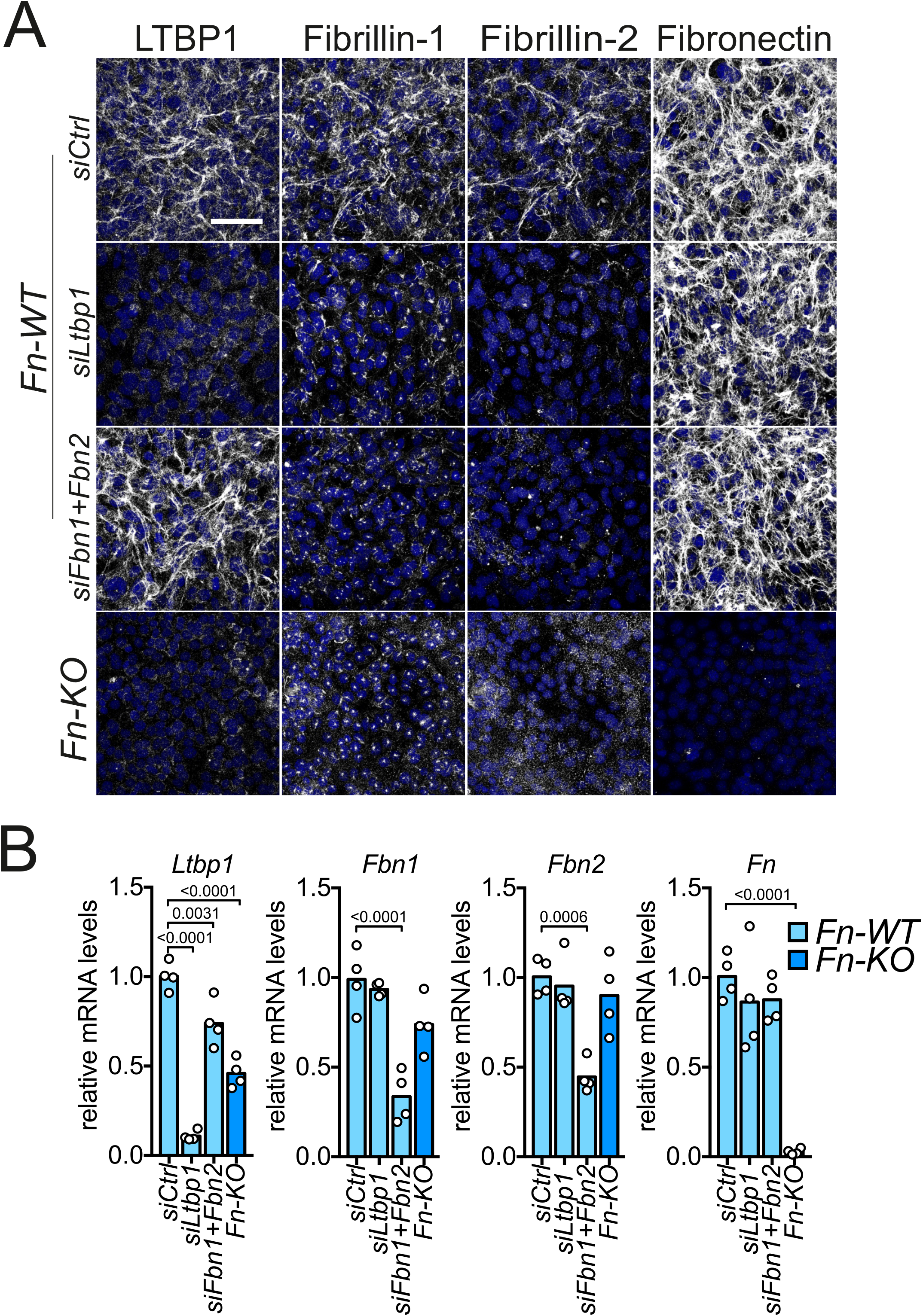
LTBP1 promotes the incorporation of fibrillins downstream of fibronectin. (**A**) Representative confocal pictures of wild-type (*Fn-WT*) and fibronectin-deficient (*Fn-KO*) MEFs cultures stained with antibodies against LTBP1, fibrillin-1, fibrillin-2 and fibronectin. Wild-type MEFs were transfected with the indicated siRNAs while fibronectin-deficient cells were left untreated. MEFs were grown for five days in medium supplemented with 10% fibronectin-depleted serum. (**B**) qPCR for *Ltbp1, Fbn1, Fbn2* and *Fn* in wild-type and fibronectin deficient MEFs transfected with the siRNAs as indicated. Data are derived from two independent experiments with two biological replicates each. 1-way ANOVA test, multiple comparisons. Significant *p*-values are reported in the plots. Scale bar: 100 μm.

### LTBP1 promotes fibrillin-1 and −2 assembly in mouse osteoblasts

Since our experiments suggest that LTBP1 may be important for the assembly of fibrillins in MEF cultures, we wanted to verify this finding in other cell types. Therefore, we generated LTBP1-deficient MC3T3-E1 calvaria osteoblasts, a murine cell line that represents a convenient system to study the hierarchical organization of microfibril-associated proteins (31). Targeting exon 7 of *Ltbp1* using CRISPR/Cas technology resulted in the generation of a cell line with abrogated expression, secretion and incorporation of LTBP1 (Figure 2A-C). By immunofluorescence analysis, we found that the linear deposition of both fibrillin-1 and fibrillin-2 was affected by LTBP1 deficiency when compared to control cells. More specifically, LTBP1-deficient cells formed fewer, shorter, fragmented fibrillin-1 microfibrils and failed almost completely to incorporate fibrillin-2 into the ECM, while fibronectin was not affected (Figure 2A). qPCR analysis revealed that fibrillin-1 expression was significantly increased in LTBP1-deficient cells while fibrillin-2 was expressed at similar levels as in wildtype cells (Figure 2B). By immunoblotting of conditioned media of LTBP1-deficient cells under non-reducing conditions we found that fibrillin-1 was secreted at higher and fibrillin-2 at slightly lower amounts (Figure 2C). To further validate our results, we transfected wildtype MC3T3-E1 with two independent siRNAs targeting LTBP1 and found that successful silencing resulted in loss of fibrillin microfibrils from the ECM, while fibronectin assembly was not grossly affected. Moreover, LTBP1 was still incorporated into the ECM of cells transfected with siRNAs against both fibrillins (Supplementary Figure 2). We also obtained similar results by transfecting primary murine calvaria osteoblasts and primary dermal fibroblasts with the same siRNAs (data not shown).

**Figure 2.**
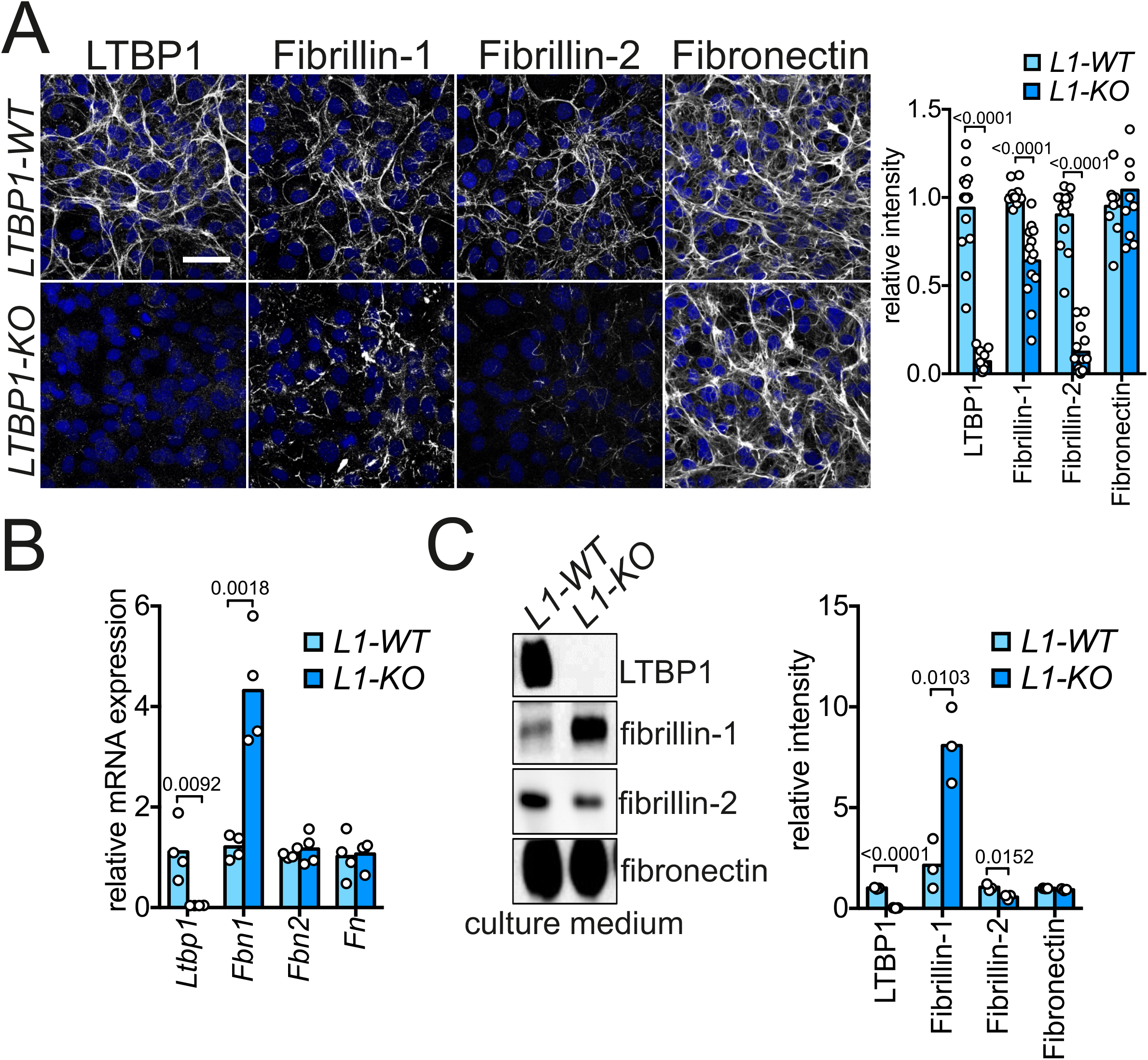
LTBP1 is required for the incorporation of fibrillins into the ECM of MC3T3-E1 osteoblasts. (**A**) Representative confocal microscopy pictures of wild-type and LTBP1-deficent MC3T3-E1 cell cultures stained with antibodies against LTBP1, fibrillin-1, fibrillin-2 and fibronectin. Note that in LTBP1-deficient cell cultures fibrillin-1 and fibrillin-2 incorporation is strongly impaired while fibronectin assembly is comparable to wild-type cultures. Quantification of images as in A. The mean fluorescence intensity was measured from confocal *z*-stack images. Data were collected from three independent experiments. (**B**) qPCR analysis for *Ltbp1, Fbn1, Fbn2* and *Fn* in wild-type and LTBP1-deficient cells. Data are derived from two independent experiments with two biological replicates each. Multiple t-tests. Significant *p*-values are reported in the plots. (**C**) Immunoblot analysis of serum-free conditioned media obtained from wild-type (*L1*-WT) and LTBP1-deficient (*L1*-KO) MC3T3-E1 osteoblasts. Samples were collected after 48 hours, concentrated five times, separated by electrophoresis on an agarose/acrylamide composite gel under non-reducing conditions and immunoblotted with the indicated antibodies. Thyroglobulin was used as a molecular weight standard (not shown). Bands represent monomeric LTBP1 and dimers of fibrillin-1, fibrillin-2 and fibronectin. Densitometric quantification of immunoblots is shown on the right. Data are from three independent experiments. Multiple t-tests were used for the statistical analysis and significant *p*-values are reported in the plots. Scale bar: 100 μm.

Altogether, our findings show that LTBP1 has an important role for the linear deposition of fibrillin microfibrils in the ECM.

### Collagens are incorporated into LTBP1-deficient ECM

Next, we asked if other ECM networks than fibrillin microfibrils are affected by LTBP1 deficiency. Therefore, we grew control and LTBP1-deficient MC3T3-E1 osteoblasts in the presence of ascorbate for 5 days and stained assembled collagen networks with antibodies against collagen I, a fibrillar collagen, collagen VI, a microfibrillar collagen, and collagen XII, a FACIT collagen. Immunofluorescence analysis showed that LTBP1-deficient cells assembled collagen networks comparable to those of control cells and semi-quantitative analysis of immunofluorescence staining revealed a more abundant collagen XII deposition (Figure 3A). However, mRNA levels of the three collagens were not affected (Figure 3B) and immunoblot analysis showed similar protein levels in cell lysates of wild-type and LTBP1 - deficient cells (Figure 3C), indicating that LTBP1 is not required for the incorporation of collagens into the ECM.

**Figure 3.**
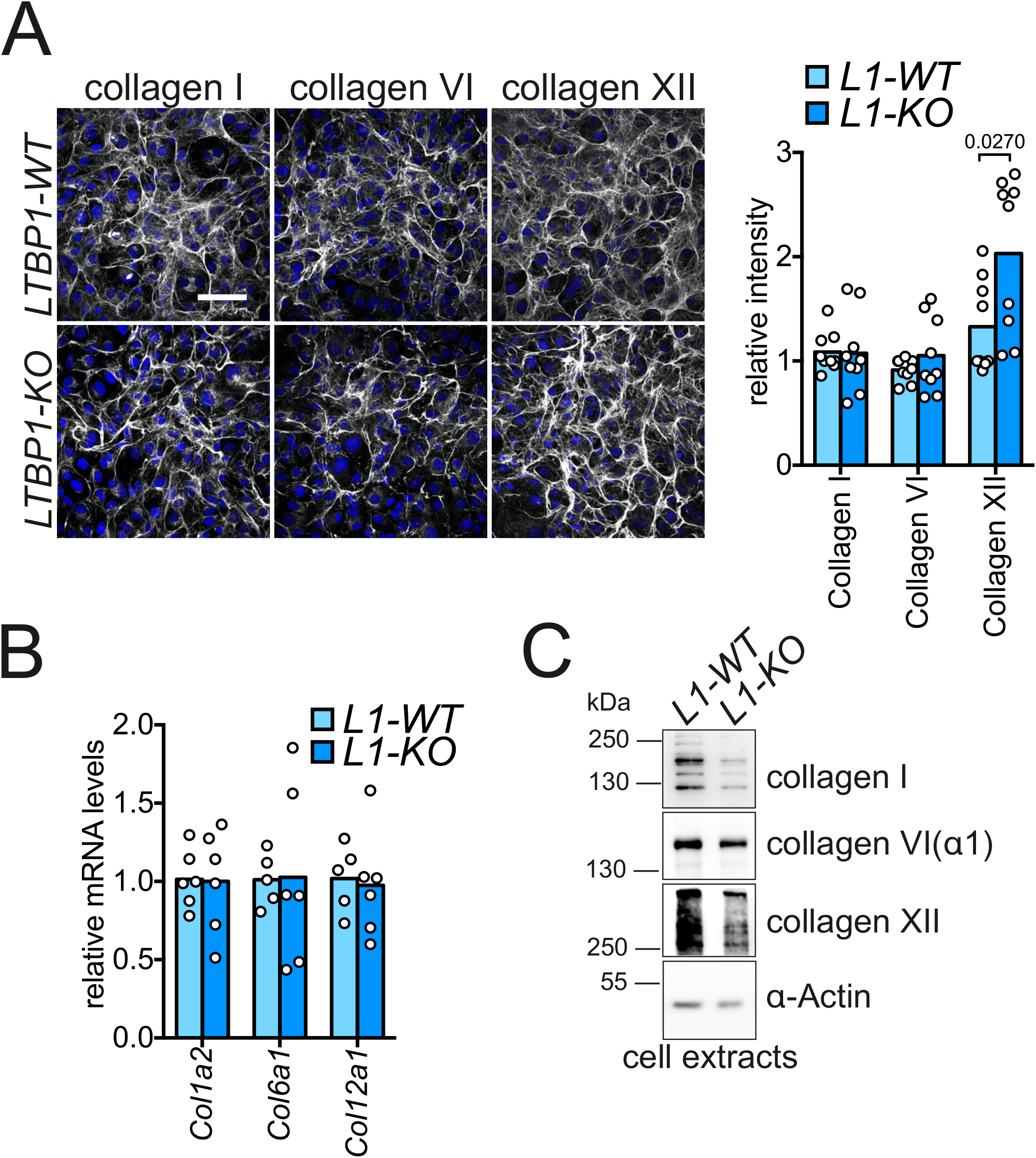
LTBP1 is dispensable for incorporation of collagens into the ECM of MC3T3-E1 osteoblasts. (**A**) Confocal microscopy images of wild-type and LTBP1-deficient cell cultures stained with antibodies against collagen I, the ⍰1 chain of collagen VI and collagen XII. The graph on the right shows quantifications of the mean fluorescence intensity as in A. The fluorescence intensity was measured from confocal *z*-stack images. Data are from three independent experiments. (**B**) qPCR for *Col1a2, Col6a1* and *Col12a1* genes. Data are derived from two independent experiments with two biological replicates each. (**C**) Immunoblot analysis of cell lysates of wild-type (*L1*-WT) and LTBP1-deficient (*L1*-KO) cells using the indicated antibodies. This experiment was performed in duplicate with similar results. Multiple t-tests were used for the statistical analysis. Significant *p*-values are reported in the plots. Scale bar: 100 μm.

### LTBP1 controls fibrillin-1 and fibrillin-2 incorporation independently from TGFβ signaling

Given the role of LTBP1 for secreting and sequestering TGFβ growth factors (9), we sought to investigate whether the impaired assembly of fibrillin-1 and −2 in LTBP1-deficient cells may be due to dysregulated TGFβ signaling. First, we examined phosphorylation levels of SMAD3 in wild-type and LTBP1-deficient MC3T3-E1 cells under basal conditions, after TGFβ1 stimulation or after treatment with the TGFβ receptor inhibitor SB431542 for 1 hour. Both treatments have been shown to be functional in MC3T3-E1 cells (32). pSMAD3 signals were not detected in wild-type and LTBP1 deficient cells, neither at basal conditions nor after treatment with the receptor inhibitor, while both cell lines showed similar levels of pSMAD3 after a 1-hour stimulation with TGF□1 (Figure 4A). qPCR analysis for two TGF□ target genes revealed an increased level of *Serpine1* (also known as *PAI-1*) and a slightly reduced level of *Col1a2* mRNA in LTBP1 deficient cells (Figure 4B). TGF□s are synthesized as small latent complexes (SLC) containing the growth factor and its cognate propeptide, the latency-associated peptide (LAP). Moreover, TGF□ can be secreted as part of the large latent complex (LLC) with the LAP covalently bound to LTBPs. Immunoblot analysis under nonreducing conditions revealed that in LTBP1-deficient cells the TGF□1 LAP was still secreted as part of the LLC (Figure 4C), suggesting that another LTBP protein is compensating for the absence of LTBP1. Altogether, these results show that the loss of LTBP1 in MC3T3-E1 cells does not result in significant changes in TGF□ signaling or secretion. Furthermore, to explore the role of TGF□ signaling in microfibril assembly, we treated both wild-type and LTBP1-deficient MC3T3-E1 osteoblasts with TGF□1 at a concentration of 2.5 ng/ml and supplied fresh medium and TGF□1 every other day. After four days, both wild-type and LTBP1-deficient cells showed a similar morphological change displaying a more spindle-shape phenotype, while no morphological difference was observed between the two groups upon DMSO or SB431542 treatment (Supplementary Figure 3). In wild-type cells, inhibition of TGF□ signaling had mild effects on fibrillin-2, but partially impaired fibrillin-1 assembly. On the other hand, stimulation with TGF□1 enhanced the deposition of thicker fibrillin-1 fibrils and prevented the incorporation of fibrillin-2 into the ECM. In LTBP1-deficient cells, neither inhibition or stimulation of TGF□ signaling had significant effects on fibrillin assembly, except for an increased accumulation of thicker fibrillin-1 fibrils that, however, remain fragmented and less developed than in wild-type cells (Figure 4D). qPCR analysis (Figure 4E) revealed that *Serpine1* and *Col1a2* mRNA levels were both similarly increased after TGF□ treatment in wild-type and LTBP-1 deficient cells. We confirmed that expression of *Fbn1* was increased in LTBP1-deficient cells under basal conditions (Figure 4E). Moreover, we found that after TGF□1 stimulation *Fbn1* mRNA levels were upregulated in both cell lines, but the upregulation was significantly stronger in LTBP1 deficient cells (2-way ANOVA, *p<0.0001*). In accordance with the immunofluorescence results, *Fbn2* expression was significantly reduced upon TGF□1 treatment in both cell lines (2-way ANOVA, *p<0.0001*), indicating that the phenotypical switch induced by TGF□1 stimulation in MC3T3-E1 cells was complemented by the loss of fibrillin-2 expression. Therefore, we conclude that TGF□ signaling has opposite effects on the expression of fibrillin-1 and fibrillin-2, and that thus LTBP1 promotes fibrillin microfibril assembly independently from its function as a gatekeeper of TGF□. To further rule out that beside TGF□s other molecules are released by LTBP1-deficient cells that inhibit the assembly of fibrillin microfibrils, we grew wild-type cells for five days in conditioned medium either derived from wild-type or LTBP1-deficient cells and assessed microfibril formation. Importantly, we found that incorporation of both fibrillins was promoted when wild-type cells were treated with medium derived from LTBP1-deficient cells, suggesting that secreted factors are not responsible for the reduction of fibrillin incorporation in the ECM of LTBP1-deficient cells (Supplementary figure 4).

**Figure 4.**
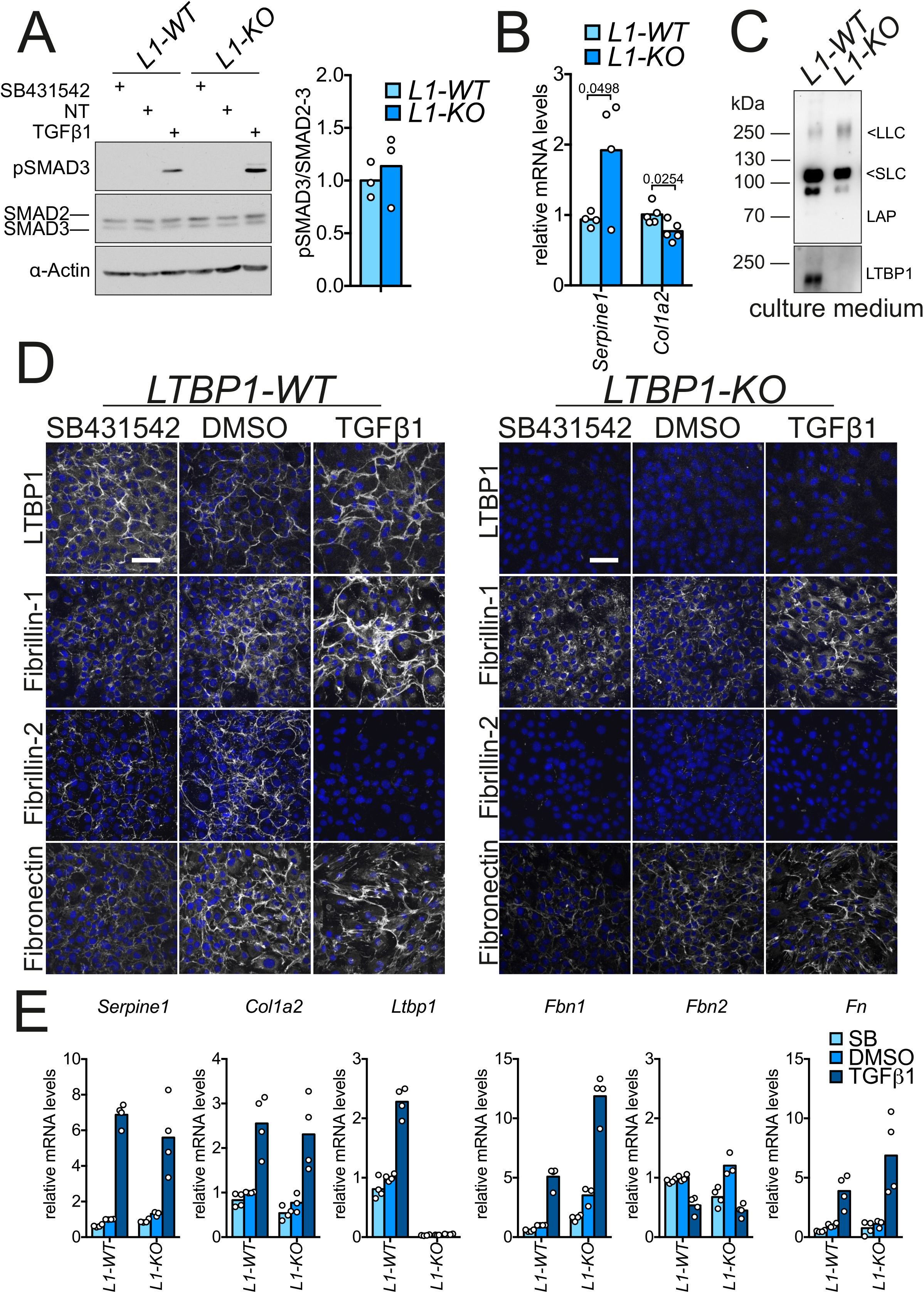
LTBP1 controls fibrillin-2 incorporation independently of TGFβ signaling. (**A**) Immunoblot for pSMAD3, total SMAD2/3 and β-actin in wild-type (*L1*-WT) and LTBP1-deficient (*L1*-KO) MC3T3-E1 cells. Cells were serum-starved overnight and treated for 1 hour with 5 ng/ml TGFβ1, 5 μM SB431542 or not treated (NT). Phosphorylated SMAD3 was only detected upon TGFβ1 stimulation. Densitometric quantification of an immunoblot as in A is shown on the right. *n*=3. (**B**) qPCR analysis of TGFβ target genes *Serpine-1* and *Col1a2* in wild-type (*L1*-WT) and LTBP1-deficient (*L1*-KO) cells. (**C**) Conditioned media of wildtype (*L1*-WT) and LTBP1-deficient (*L1*-KO) cells were separated by electrophoresis on a 4-12% gradient gel under non-reducing conditions and immunoblotted with the indicated antibodies. (**D**) Confocal microscopy images of wild-type and LTBP1-deficient MC3T3-E1 treated for 4 days with DMSO, 2.5 ng/ml TGFβ1 or 5 μM SB431542 and stained with the indicated antibodies. (**E**) qPCR for *Serpine1, Col1a2, Ltbp1, Fbn1, Fbn2* and *Fn* from cDNA generated from cells treated as in D. Data are derived from two independent experiments. NT: non-treated. Scale bars: 100 μm.

### Recombinant LTBP1S promotes fibrillin-1 and −2 incorporation with different efficiency

Next, to test if LTBP1 may have a direct role for fibrillin-1 and −2 incorporation, we treated LTBP1-deficient cells with purified LTBP1S, the better characterized of the two LTBP1 isoforms. First, we verified that the recombinant protein could be efficiently incorporated in the ECM of LTBP1-deficient cells (Figure 5A). The addition of LTBP1S to the cell culture medium promoted the incorporation of fibrillin-2 with high efficiency, while the effect on the linear deposition of fibrillin-1 was much less pronounced (Figure 5B). We also observed that LTBP1-deficient cells were able to incorporate recombinant LTBP1S into the ECM even after siRNA-mediated double knockdown of fibrillin-1 and fibrillin-2, confirming that LTBP1 does not strictly depend on fibrillins for its linear deposition (Figure 5A). qPCR analysis revealed that LTBP1S supplementation did not significantly affect the mRNA levels of *Serpine1, Fbn1* and *Fbn2* (Figure 5C).

**Figure 5.**
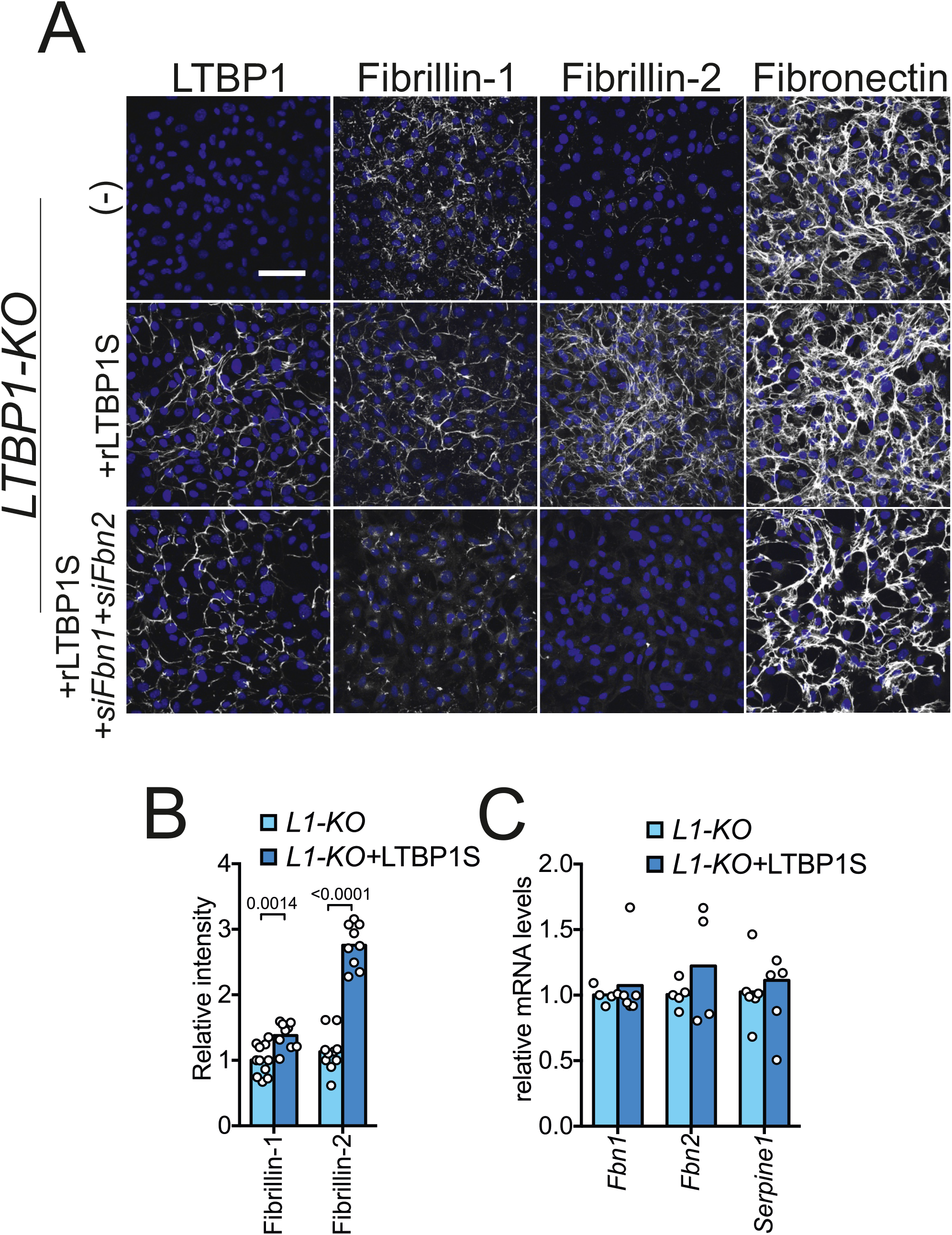
Effects of recombinant LTBP1S on the incorporation of fibrillins in the ECM of LTBP1-deficient cells. (**A**) Representative confocal microscopy images of LTBP1-deficient MC3T3-E1 cultures that were left untreated or transfected with siRNAs specific for fibrillin-1 and 2 (*siFbn1+siFbn2*). The next day cells were treated with fresh medium with 15 nM recombinant LTBP1S (+rLTBP1) or left untreated (-) for four more days. Supplementation of recombinant LTBP1S strongly promoted the assembly of fibrillin-2, while having only a modest effect on fibrillin-1 microfibrils. siRNA-mediated knockdown of both fibrillins did not prevent the incorporation of LTBP1S and confirmed the specificity of the antibodies. (**B**) Quantification of the mean fluorescence intensity as in A for fibrillin-1 and fibrillin-2. The fluorescence intensity was measured from confocal *z*-stack images. The mean fluorescence intensity of knockout cells was set as 1. Data are from two independent experiments. (**C**) qPCR analysis showed that the treatment with soluble LTBP1S did not grossly affect the mRNA levels of the TGFβ target gene *Serpine1* nor of *Fbn1* and *Fbn2*. Data are from two independent experiments. Multiple t-tests were used for the statistical analysis. Significant *p*-values are reported in the plots. Scale bar: 100 μm

### LTBP1S and LTBP1L have distinct activities in ECM assembly

Our data using recombinant LTBP1S imply that the short form of LTBP1 have different effects on the incorporation of the two fibrillins. LTBP1 can be synthesized as a short form (LTBP1S) or as an N-terminally extended long form (LTBP1L) (Figure 6A). We hypothesized that the two forms may have different properties within the ECM. Using RT-PCR analysis we found that both forms are endogenously expressed by wild-type MC3T3-E1 cells (not shown). Therefore, to test our hypothesis, we generated two LTBP1-deficient MC3T3-E1 cell lines stably expressing either LTBP1S or LTBP1L. Strikingly, and in accordance to previous findings (12), we observed by immunofluorescence that LTBP1L was incorporated into the ECM with a higher efficiency than LTBP1S (Figure 6B,C), and qPCR analysis showed similar expression levels for the two constructs (Figure 6D). Interestingly, LTBP1L was also much more efficient in promoting the assembly of fibrillins, and this different activity was more prominent for the linear deposition of fibrillin-1 than for that of fibrillin-2 (Figure 6B,C). The increased impact on fibrillin-1 deposition may be due to the much higher tendency of LTBP1L to be incorporated into the ECM when compared to LTBP1S or to an intrinsic functional difference between the two isoforms. We also tried to recapitulate a similar effect by treating LTBP1-deficient cells with increasing concentrations of recombinant LTBP1S. However, even at a concentration of 100 nM, the short isoform was not able to promote the linear deposition of fibrillin-1 as efficiently as the long form (not shown). qPCR analysis revealed that LTBP1S and LTBP1L cells expressed *Fbn2* at a significantly higher level than LTBP1-deficient cells, while only LTBP1L caused a significant increase in *Fbn1* expression. Finally, in agreement with the finding that LTBP1-deficient cells showed slightly increased *Serpine1* levels compared to wild-type cells (see Figure 4B), both LTBP1 isoforms slightly reduced its expression (Figure 6D). Altogether these results show that the two LTBP1 isoforms are incorporated with different efficiency in the ECM where they may have distinct roles for the assembly of fibrillin microfibrils.

**Figure 6.**
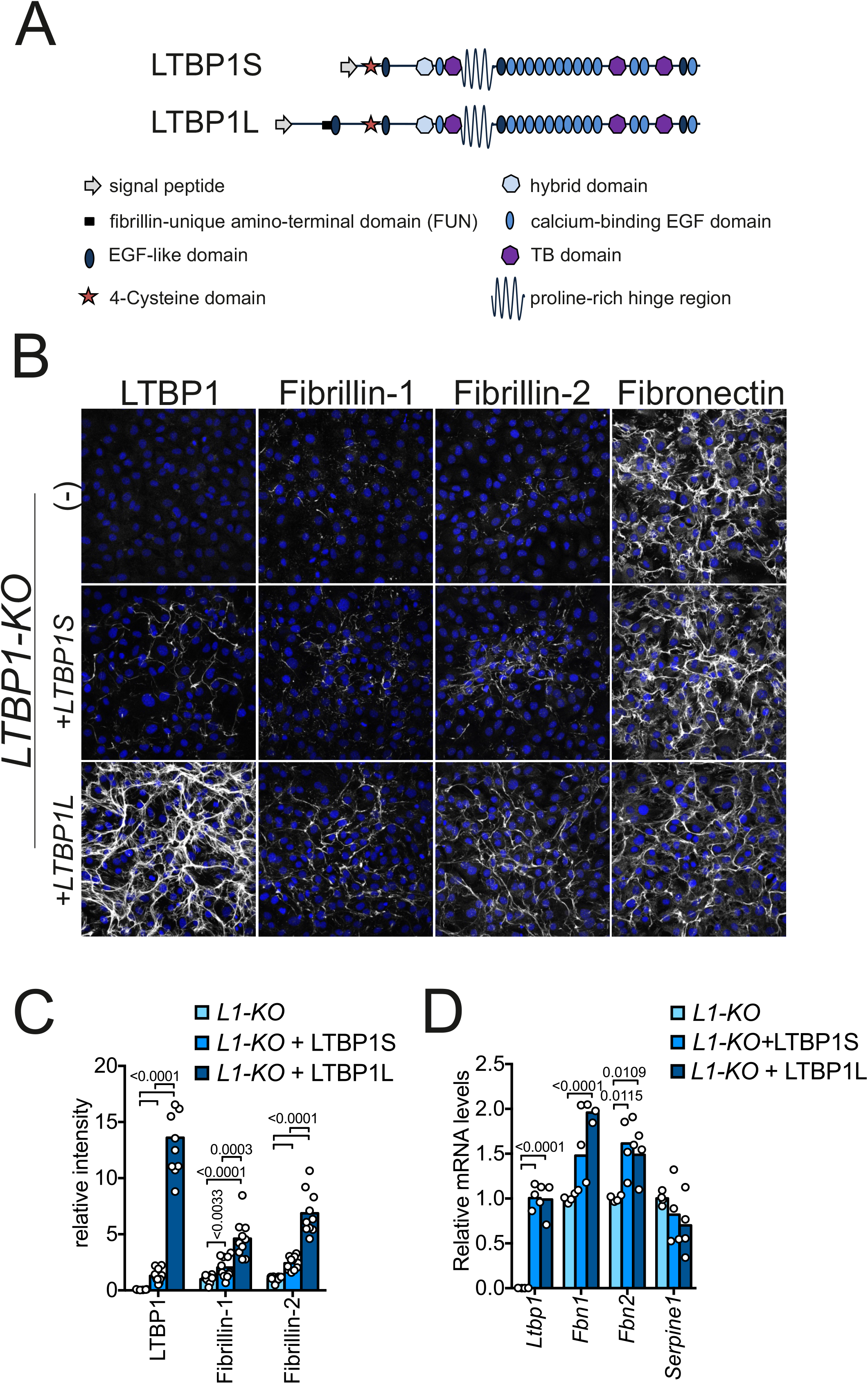
LTBP1L incorporate into the ECM and promote the linear deposition of fibrillin microfibrils more efficiently than LTBP1S. (**A**) Schematic representation of LTBP1S and LTBP1L domain organization (adapted from (9)). (**B**) Representative confocal microscopy images of non-transfected LTBP1-deficent MC3T3-E1 cultures (-) and of LTBP1-deficient cells transfected with constructs coding for the two LTBP1 isoforms (*+LTBP1S* and +*LTBP1L*). Cells were grown for five days on glass coverslips before fixation and stained with the indicated antibodies. (**C**) Quantification of the mean fluorescence intensity as in A for LTBP1, fibrillin-1 and fibrillin-2. The fluorescence intensity was measured from confocal *z*-stack images. Data are from two independent experiments. For LTBP1, the mean fluorescence intensity of LTBP1S-epressing cells was set as 1, while for fibrillin-1 and −2 values from the knockout cells were used as reference. (**D**) qPCR of *Ltbp1, Fbn1, Fbn2* and *Serpine1* in the three cell lines. Multiple t-tests were used for statistical analysis. Significant *p*-values are reported in the plots. Scale bar: 100 μm

## Discussion

LTBP1 was originally discovered during the purification of TGF□ from human platelets (33). It subsequently became obvious that LTBP1 — similar to other LTBPs — is related to fibrillin-microfibrils, both because of the structural homology and their direct association with fibrillins within tissues (8,34,35). Despite its structural similarity to the other LTBPs and their shared targeted localization to fibrillin microfibrils, LTBP1 has a unique ability to assemble in the ECM. Indeed, while it has been shown that LTBP2, LTBP3 and LTBP4 strictly depend on fibrillin-1 for their incorporation into the ECM, LTBP1 requires fibronectin (29,30,35). Notably, mouse embryonic fibroblasts lacking both fibrillins as well as calvarial osteoblasts derived from fibrillin-2 knockout mice could still form LTBP1 fibrils (29,36). Therefore, LTBP1 assembly appears not to depend on fibrillin microfibrils and our data support this concept. However, the molecular determinants that confer LTBP1 with this specificity remain unknown. Like LTBP1, fibrillin-1 and −2 were also found to depend on fibronectin for their assembly into the ECM (27,28), but the precise interdependent relationships between these ECM components — fibronectin, fibrillins and LTBP1 — remained unsolved. Our results propose a new hierarchical model in which LTBP1 promotes the linear deposition of fibrillin microfibrils. This idea is also consistent with previous studies showing that in cultures of osteoblasts and osteosarcoma cells, LTBP1 seems to associate first with fibronectin and later with fibrillin microfibrils (30,37). LTBP1 binds directly to the N-terminal region of both fibrillins via its C-terminal region (36,38). On the other hand, it is unclear how LTBP1 associates with fibronectin fibrils. Indeed, while a direct binding between fibronectin and an LTBP1 peptide spanning residues 21-629 was initially reported (39), a subsequent study failed to confirm such an interaction. The authors instead suggested that the association could be mediated by heparan sulphate proteoglycans. Consistently, heparin treatment resulted in the lack of LTBP1 incorporation into the ECM of fetal rat calvaria cells, while leaving the fibronectin network unaffected (40). Interestingly, a similar role for heparan sulphate proteoglycans has been demonstrated in the assembly of fibrillin-1 and −2 (41). At present, the specific heparan sulphate proteoglycan mediating such effects are unknown, it is however tempting to speculate that LTBP1 may support fibrillin assembly at least in part by modulating heparan sulphate proteoglycans function/structure in the extracellular milieu.

LTBP1 is synthesized in both long and short forms by the use of unique translation initiation sites. LTBP1L-deficient mice die shortly after birth because of the incomplete development of the cardiac outflow tract, a defect that has been linked to decreased TGF□ signaling (42,43), while mice lacking only the short form show mild craniofacial and skeletal abnormalities (42,44,45). However, it is not clear if the different phenotypes depend on distinct functions of the two isoforms, as the biological significance of the long and short forms is still unresolved (9). Our data, in agreement with a previous study (12), show that the two isoforms are endowed with distinct properties. Indeed, we found that when expressed in LTBP1-deficient MC3T3-E1 cells, LTBP1L is incorporated more efficiently into the ECM than LTBP1S, and complemented by an increased capacity to promote the linear deposition of fibrillins. LTBP1L in addition to the sequence of the short form, contains an N-terminal 346 amino acid FUN (Fibrillin Unique N-terminal) domain paired with an EGF-like domain. Interestingly, this domain pair is found only in fibrillins, LTBP1L and LTBP2 (46). The precise function of the FUN domain is not known, but mutations mapping to the FUN domain of fibrillin-1 have been reported in patients affected by Marfan syndrome (47,48). Detailed studies are therefore needed to pinpoint more precisely the functions of the FUN domain and to explore if FUN is the reason for the different behavior of the two LTBP1 isoforms.

The recent identification of novel disease-causing variants of *LTBP1* adds further insights into its function. Indeed, the core clinical features of the *LTBP1*-related syndrome, including cutis laxa, craniosynostosis, characteristic craniofacial defects, short stature, brachydactyly and only mild cardiovascular phenotypes, are partially overlapping with the manifestations of other fibrillinopathies and point to specific multifaceted functions of LTBP1 in the homeostasis of connective tissues. Notably, although the two analyzed variants were found to have a different impact on TGF□ signaling in cultures of patient fibroblasts — namely increasing it or not affecting it— they both showed similar outcomes at the phenotypical level. Therefore, it is likely that other TGF□-independent functions may be compromised in the differently mutated LTBP1 proteins. In line with this idea, truncations in different regions of LTBP1 caused distinct defects in ECM assembly in patient fibroblasts (23). Our data show that complete loss of LTBP1 causes a significant impairment of fibrillin microfibril assembly in cell culture, while not resulting in major dysregulation of canonical TGF□ signaling. It is therefore tempting to speculate that impaired fibrillin microfibril biogenesis/proteostasis may contribute to the pathogenesis of the *LTBP1*-related syndrome. Notably, both deficiency and increased deposition of fibrillin-1 and fibrillin-2 lead to skeletal anomalies (36,49,50), and skeletal features are prominent in individuals harboring *LTBP1* pathologic variants (23). Interestingly, different fibrillin-1 mutations cause opposite skeletal phenotypes, tall stature and arachnodactyly in Marfan syndrome and short stature and brachydactyly for Weill-Marchesani syndrome. Moreover, fibrillin-2 deficiency in zebrafish has been linked to defects in notochord formation and *Ltbp1*^-/-^ zebrafish display several vertebral defects (23,51). Finally, mutations disrupting the *Fbn2* gene cause syndactyly in mice, a feature that has also been described in five of the eight LTBP1 patients described so far (23). In summary, in this study we have unveiled a novel non-TGF□-dependent function of LTBP1, that may contribute to the pathogenesis of the recently-identified *LTBP1*-related human disorder and extends our knowledge on the complex process of extracellular microfibril assembly.

## Material and Methods

### Antibodies

The polyclonal antibodies against LTBP1, fibrillin-1, fibrillin-2, the collagen VI ⍰1 chain and collagen XII were previously described (23,52-55). The polyclonal antibody against collagen I was from Cedarlane (CL50151AP) and those against pSMAD3 (C25A9) and total SMAD2/3 (D7G7) were from Cell Signaling Technology. A goat anti-LAP (TGF□1) antibody was from R&D Systems.

### Constructs

cDNA encoding for full length human LTBP1L (NM_206943; AA: 24-1721) and LTBP1S (BC144128; AA: 21-1395) were generated by gene synthesis (GeneArt^®^ Strings, Thermofisher) and used for PCR amplification. PCR products were digested with the appropriate restriction enzymes and cloned into a modified Sleeping Beauty transposon expression vector containing a BM-40 signal peptide followed by a Twin-Strep-Tag (56,57).

### Purification of recombinant LTBP1S

For protein production, the LTBP1S expression construct was co-transfected with the transposase plasmid (10:1) into HEK293T cells. After puromycin selection (3 μg/ml), cells were expanded in triple flasks and when confluency was reached, protein production was induced with doxycycline (0.5 μg/ml). Cell supernatants were harvested every 3 days, and the recombinant proteins purified via the Strep-Tactin®XT (IBA Lifescience) resin by elution with biotin containing TBS-buffer (IBA Lifescience).

### Cell culture and siRNA transfection

MC3T3-E1 cells (subclone 4) were purchased from ATCC and maintained in alpha Minimum Essential Medium supplemented with 10% FBS, 1 % penicillin/streptomycin. Fibronectin wild-type and knockout embryonic fibroblasts were a kind gift from Dr. Reinhard Fässler (58). siRNA reverse-transfections were carried out with Lipofectamine RNAiMAX (Invitrogen) according to the manufacturer’s instructions. The AllStars Negative Control siRNA and gene-specific siRNAs were purchased from Qiagen and Eurofinns, respectively. siRNA sequences used in this study were: *siLtbp1 #1*: 5’TACGCAAGTAACAGAAATCAA3’; *siLtbp1 #2:* 5’AAGGGAGAGATGTATACTTGA3’; *siFbn1:* 5’ATGGTGCTTATTAAGACCAAA3’; *siFbn2:* 5’CTCGACGAATGTCAAACCAAA3’. For ECM network formation and RNA analysis, cells were seeded on uncoated glass coverslips or directly on plastic, respectively.

### Generation of LTBP1 KO MC3T3-E1 cells and LTBP1L and LTBP1S expressing cells

Knock-out cell lines were generated using the CRISPR/Cas9 system (59). In brief, doublestranded DNA oligonucleotides that encode guide RNAs (gRNAs) against the target gene were cloned into the BbsI restriction sites of the PX459 vector. The guide sequence used for the sgRNA expression plasmids to generate the LTBP1-KO line targeting the exon 7 was: 5’TATTCCCCATGTGTATCCCG3’. MC3T3-E1 cells were transfected with the resulting vectors and selected with puromycin (1.5 μg/ml) for 4 days. Single cell clones were picked using cloning cylinders and a knock-out clones were validated by immunoblotting. The respective control cell line was generated by using the empty vector and following the same procedure. To generate LTBP1L and LTBP1S expressing cells, constructs were transfected in LTBP1-deficient MC3T3-E1 cells with the transposase plasmid (10:1) and the cells were selected for one week with puromycin at a concentration of 1,5 μg/ml.

### Immunofluorescence and confocal microscopy

For ECM network formation cells were seeded in 24-well plates on uncoated glass coverslips at a density of 8□ × □10^4^ cells/well. The next day the medium was changed and the cells were grown for 4 more days in DMEM supplemented with 10% FCS and 1 % penicillin/streptomycin. For collagen biosynthesis the medium was supplemented with ascorbate. When indicated, the medium was supplemented with 2.5 ng/ml TGFβ1 (Preprotech) or 5 μM SB431542 (Merck) and changed every 2 days. Cells were fixed for 5 minutes at −20°C in methanol/acetone, blocked with 1% FBS in PBS for 30 minutes, incubated with primary antibodies diluted in 1% FBS in PBS for 1 hour at RT, washed 3x with PBS for 10 minutes, and incubated with appropriate highly cross-adsorbed secondary antibodies conjugated to Alexa Fluor 488 or 555 (Thermo Fisher Scientific) and DAPI (0.1 μg/ml; Sigma-Aldrich) for 1 hour at RT. Samples were mounted with Dako fluorescent mounting medium (Agilent/Dako). Images were acquired by confocal microscopy using a Leica SP5 system controlled by the LAS AF 3 software (Leica Microsystems). For semi-quantitative quantification of the fluorescence intensity, at least 5 z-stacks (3-4 μm thick) were randomly acquired for each coverslip and the mean fluorescence intensity calculated with the Leica LAS AF Lite software (figures 2-3) or with ImageJ (figures 5 and 6). Images were processed with the ImageJ software.

### Real-time PCR

Cells were seeded on 12-well or 24-well plates at a density of 1.6□×□10^5^ or 8□×□10^4^ cells/well, respectively and cultured for 5 days. Total RNA was prepared with the Trizol™ reagent (Invitrogen) following the manufacturer’s protocol. RNA (1 μg per sample) was reverse transcribed using the Qiagen OmniScript RT Kit. Technical duplicates for every sample were amplified using the Takyon ROX SYBR 2X MasterMix dTTP blue (Eurogentec) in a StepOnePlusTM Real-Time PCR Detection System (Applied Biosystems). Data analysis was performed using the 2–ΔΔCt method and quantified relative to *Gapdh*. The primer sequences are provided in Supplementary Table 1.

### Immunoblot

Control and LTBP1-deficient cells were plated in 6-well plates at a density of 1.6□×□10^5^/well. After two days, cells were switched for additional 48 hours in serum-free DMEM. The conditioned media were precipitated, resuspended in 2x Laemmli sample buffer and subjected to electrophoresis in 0.5% (w/v) agarose/2.4% (w/v) polyacrylamide composite gels or in 4-12% polyacrylamide gradient gels under non reducing conditions (60). Thyroglobulin (660 kDa) (Sigma) was used as a molecular weight standard. For pSMAD3 immunoblots, cells were starved overnight in serum-free DMEM and then treated for 1 hour with 5 ng/ml TGFβ1 (Preprotech) or with 5 μM SB431542 (Merck), and intensities quantified by ImageJ. A density profile was obtained and the area under each peak was measured as the number of pixels, normalized to total SMAD2/3 and calibrated by fixed point of control (61).

### Statistical analysis

Statistical analyses were performed with the GraphPad Prism software.

## Acknowledgments

This work was supported by the Deutsche Forschungsgemeinschaft through project ID 384170921: FOR2722/B1 (MPa), FOR2722/B1 (RW), FOR2722/B2 (MK) and FOR2722/ C2 (GS).

## Author contributions

AS, MPr, VSG and BK performed the experiments. FM, SM and MK produced recombinant LTBP1S. GS, RPM and MK provided antibodies and plasmids. BB, GS, MPa, MK and RW, funding acquisition, resources and critical comments. AS designed and supervised the study, and wrote the manuscript. All the authors read and commented on the final manuscript.

---

**Supplementary Figure 1.**
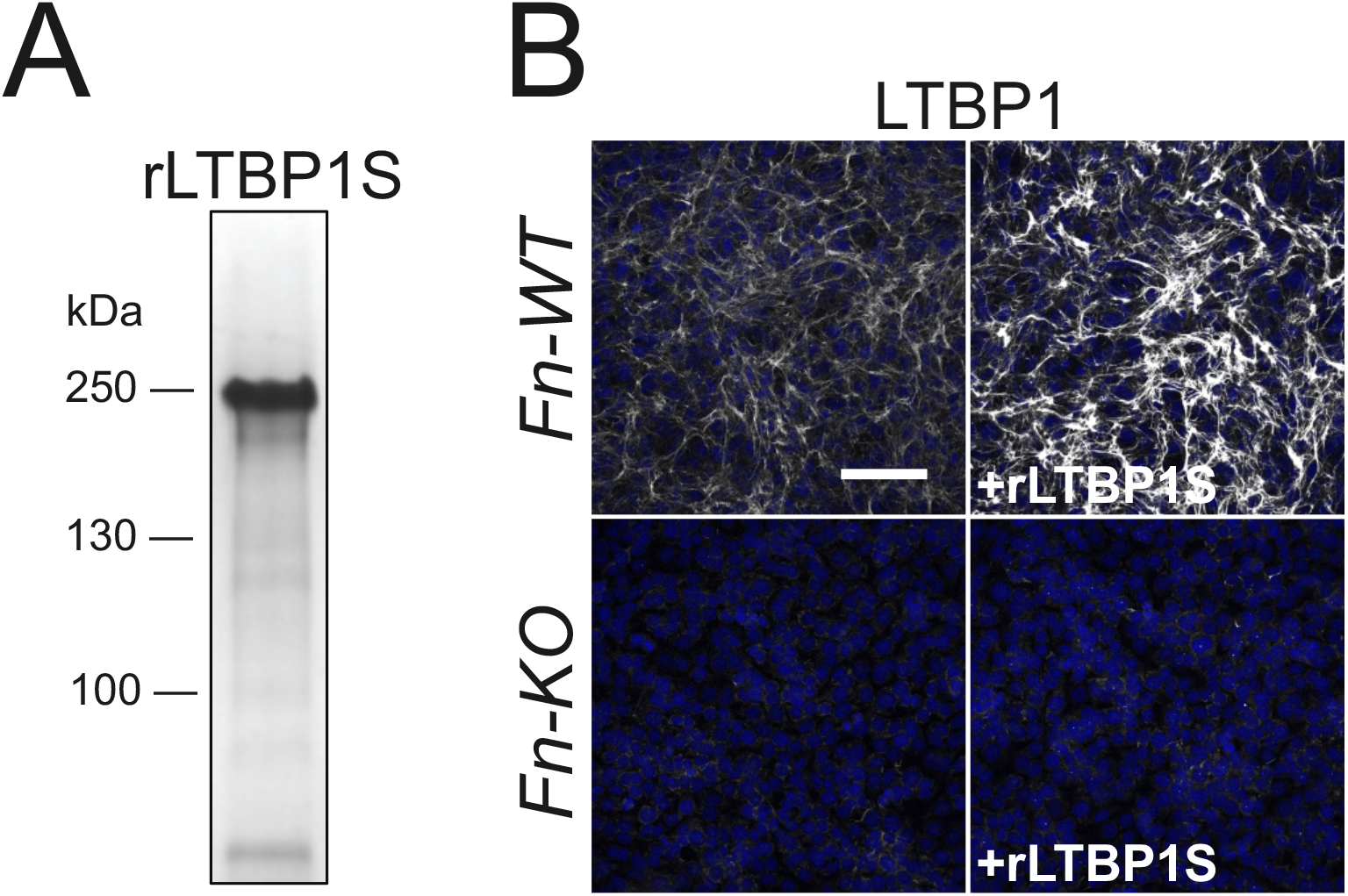
Fibronectin is required for the incorporation of LTBP1 into the extracellular matrix. (**A**) Coomassie Blue stained SDS/polyacrylamide gel electrophoresis of the recombinant human LTBP1S protein used in this study. (**B**) Wild-type and fibronectin-deficient MEFs were left untreated or treated with 15 nM recombinant LTBP1 (+rLTBP1). After five days in culture cells were fixed and stained with the antibody specific for LTBP1. Scale bars 100 μm

**Supplementary figure 2.**
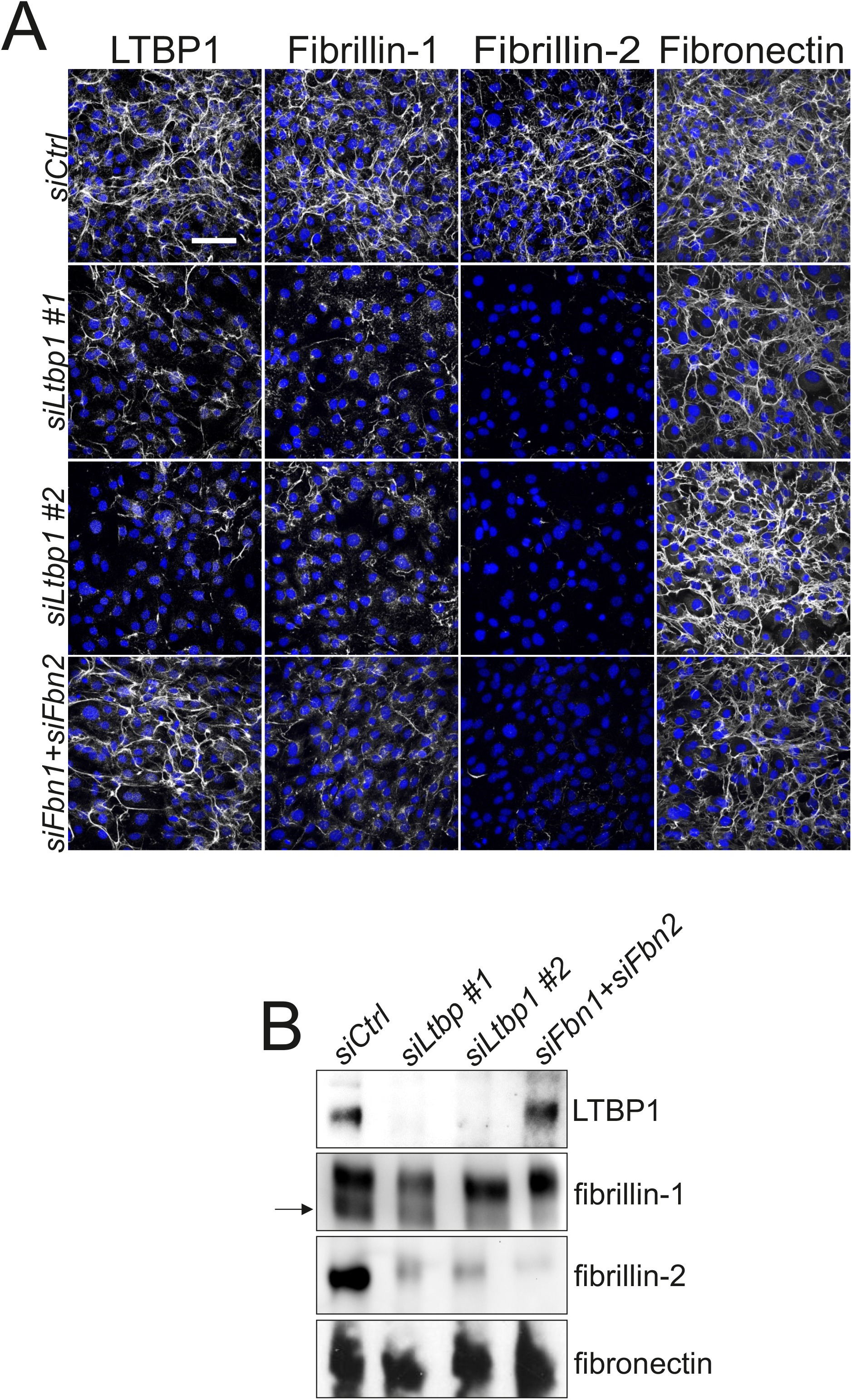
LTBP1 is required for the incorporation of fibrillins into the ECM of MC3T3-E1 osteoblasts. (**A**) siRNA mediated knockdown of LTBP1 and fibrillin-1 and −2 in MC3T3-E1 wild-type osteoblasts. After five days transfected cell cultures were fixed and stained with antibodies directed against LTBP1, fibrillin-1, fibrillin-2 and fibronectin. (**B**) Immunoblot analysis of serum-free conditioned media obtained from MC3T3-E1 osteoblasts transfected as in A. Samples were collected after 48 hours, concentrated five times, separated by electrophoresis in an agarose/acrylamide composite gel (for fibrillin-1, fibrillin-2 and fibronectin) or in a 4-12% gradient gel (for LTBP1) under non-reducing conditions and immunoblotted with the indicated antibodies. Thyroglobulin was used as a molecular weight standard (not shown). Bands represent monomeric LTBP1 and dimers of fibrillin-1, fibrillin-2 and fibronectin. The arrow points to the specific band for fibrillin-1. Scale bar: 100 μm

**Supplementary figure 3.**
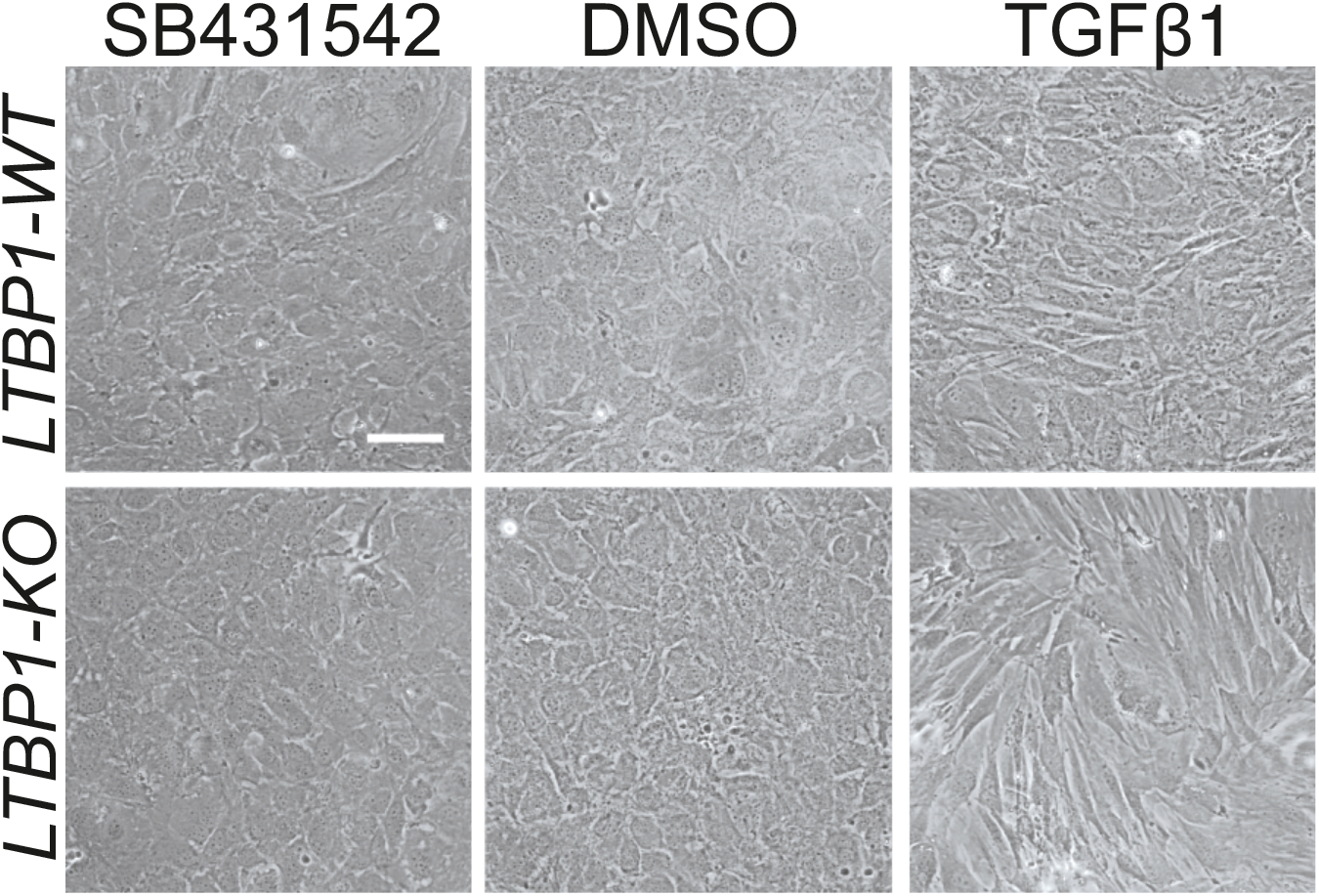
Effects of SB431542 and TGFβ1 on MC3T3-E1 cell morphology. Bright-field microscopy images of control and LTBP1-deficient MC3T3-E1 cells treated for four days with DMSO, 2.5 ng/ml TGFβ1 or 5 μM SB431542. Note the morphological shift induced by TGFβ1 in both cell lines. Scale bar: 100 μm

**Supplementary figure 4.**
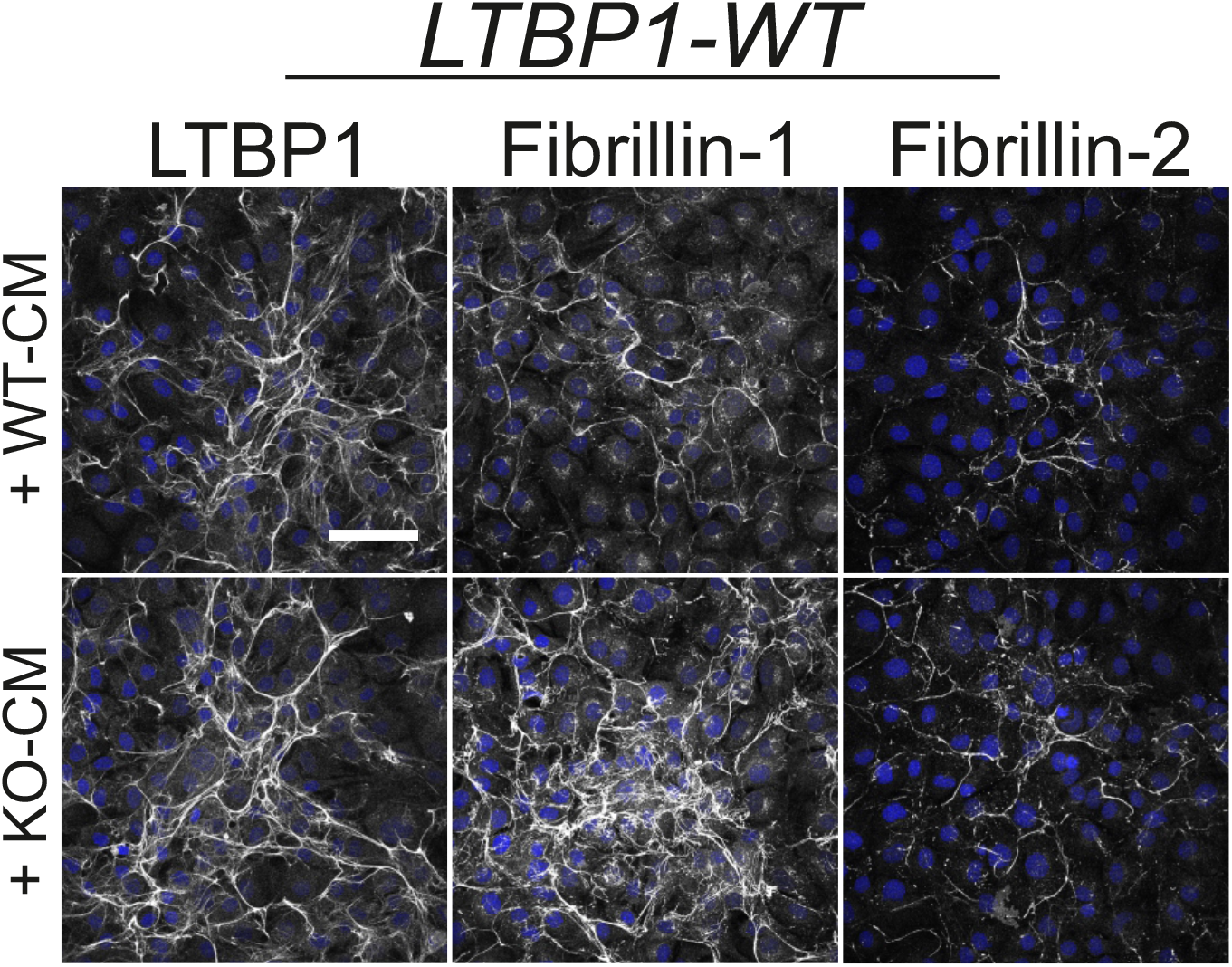
Wild-type MC3T3-E1 cells were grown for five days with 7-days conditioned media derived from culture of either control (+WT-CM) or LTBP1-deficient cells (+KO-CM) and stained with the indicated antibodies. In agreement with the increased secretion level of fibrillin-1 in LTBP1-deficient cells more abundant fibrillin-1 microfibrils were deposited by the cells treated with the knockout medium. Scale bar: 100 μm

